# Mutually Beneficial Symbiosis Between Human and Gut-Dominant *Bacteroides* Species Through Bacterial Assimilation of Host Mucosubstances

**DOI:** 10.1101/2020.08.21.262261

**Authors:** Masahiro Sato, Kanta Kajikawa, Tomoya Kumon, Daisuke Watanabe, Ryuichi Takase, Wataru Hashimoto

## Abstract

The composition of gut microbiota is influenced by the quantity and type of nutrients in host. Even with some *Bacteroides* species being categorized as pathogens, *Bacteroides* is one of the most dominant gut bacteria. Here we indicate the physiological determinants of the species of *Bacteroides* for being dominant in human gut microbiota. Each of the host extracellular mucosubstances including glycosaminoglycans (GAGs) and mucin has grown human gut microbiota. In spite of the differences among initial microbiota profiles, *Bacteroides* species dominated the community when GAG (e.g., chondroitin sulfate or hyaluronan) was used as a sole carbon source. In fact, GAGs and the *Bacteroides* genes which are vital for the degradation of GAGs were commonly detected in human feces. Mucin has encouraged the growth of *Bacteroides* and several other genera. A comprehensive analysis on the degradation and assimilation of mucosubstances by the genus *Bacteroides* using around 30 species has shown that most species degrade and assimilate GAGs and mucin, showing that *Bacteroides* species can survive even in the undernutrition condition including the fasting state. In the assimilation of GAG or mucin, *Bacteroides* species significantly secreted essential amino acids, γ-amino butyrate (GABA), and/or short-chain fatty acids which are needed for human health. This is the first report as regards mutually beneficial interaction between human and *Bacteroides* species via bacterial assimilation of host mucosubstances and secretion of metabolites for host health promotion.

**Significance:** The genus *Bacteroides* is one of the most dominant gut bacteria, although its beneficial effects on human health have not been well understood. Here, we show modes of action in human-*Bacteroides* interrelationship. Mucosubstances including GAGs and mucin secreted by human host are abundant in gut for microbiota to grow well. *Bacteroides* species are dominant in the community in the presence of GAGs, and provide human host with a considerable amount of essential amino acids, γ-amino butyrate, and short-chain fatty acids produced from mucosubstances. These results postulate mutually beneficial symbiosis system between human and *Bacteroides* through bacterial assimilation of host mucosubstances and secretion of metabolites for human body and mental health promotion even in the undernutrition condition including the fasting state.

## Introduction

In the human gut, there are 100 trillion bacteria communicating with each other and forming a complex microbiota (1). The gut microbiota’s composition is influenced by different host diets and metabolizable nutrition for bacteria (2). Contrarily, gut bacteria colonize mucosubstances including mucus layer mucin or extracellular matrix glycosaminoglycan (GAG) made by the host to preserve the intestine (3). Mucin is among the principal components of the mucus layer of the intestine and is comprised of the main chain protein and side chain sugars (e.g., *N*-acetylglucosamine (GlcNAc), *N*-acetylgalactosamine (GalNAc), galactose, fucose, sialic acid) (4). GAG is comprised of a two-sugar component-repeated structure, uronate like iduronic acid or glucuronic acid (GlcUA), and amino sugar including GlcNAc or GalNAc (5). In amino sugars, both hydroxyl and amino groups are highly sulfated. GAG is categorized into chondroitin sulfate (CS), hyaluronic acid (HA), heparin (HP), heparan sulfate, dermatan sulfate, and keratan sulfate based on constituent sugar, glycoside bond manner, and the level of sulfated groups (5).

Some human gut bacteria, for example, *Bacteroides, Clostridium, Enterococcus*, and *Streptococcus* genera, break down GAGs that are supplied by the host intestine which are not dependent on intake of meals (6–9). Our previous studies have exhibited molecular machineries of depolymerization, import, degradation, and metabolism of GAGs in the genus *Streptococcus* (10, 11) (*SI Appendix*, Fig. S1). Mainly among these gut bacteria, *Bacteroides* species assimilate polysaccharides that are supplied by the host and colonize the colon intestinal mucus layer as the largest group in humans in spite of race (12–14), even though the genus *Bacteroides* is often classified as an opportunistic pathogen (15). *Bacteroides thetaiotaomicron* assimilates both mucin *O*-binding polysaccharides and GAGs including CS, HA, and HP (16–18).

On the other hand, short-chain fatty acids which are made by some gut bacteria positively influence the host physiology, for example (19), (i) improve the gut environment by decreasing the pH, (ii) adjust the host immune system, (iii) be the energy source of the host, (iv) improve insulin resistance, and (v) decrease the risk of a heart attack. Some species of *Bacteroides* can create short-chain fatty acids (20), indicating their function as next-generation probiotics (15, 21, 22). For one to know the reason why humans allow *Bacteroides* species to be dominant in gut microbiota and why humans do not repress the bacterial species, this article deals with the mechanism of symbiosis between humans and gut-dominant genus *Bacteroides* through comprehensive analyses to explore *Bacteroides* species capability to degrade as well as assimilate human mucosubstances (GAGs and mucin) and to dictate bacterial metabolites during the human secretion assimilation.

## Results

### GAG Is Available to Human Gut Microbiota

Extracellular matrices under the mucus layer which is comprised of mucin (23) are generally composed of GAGs, including CS, HA, and HP. Since the mucus layer hinders bacteria from invading the host cells (24), these bacteria can hardly utilize GAGs. On the contrary, human feces have components from epithelial cell shedding (25), indicating that GAGs that are from the intestinal tracts are available to the gut microbiota. Therefore, the concentration of GAGs, especially those in sulfated forms, in feces was measured using the 1,9-dimethyl-methylene blue (DMMB) method (26). The contents of GAGs in independent feces from three volunteers A to C (two twenties and one fifties) were as follows: volunteer A, 0.098% (w/w); volunteer B, 0.112%; and volunteer C, 0.033%. These values are similar to those of GAGs that are part of bacterial media, indicating that there are plenty GAGs in the human gut for microbiota to grow well. In fact, *B. thetaiotaomicron* has shown a remarkable growth in the minimal medium with the use of 0.02% (w/v) GAG (HA) as a sole carbon source (*SI Appendix*, Fig. S2).

### *Bacteroides* Species Are Dominant in the Community in the Presence of GAG

To explore the effects of carbon sources on human gut microbiota, fecal samples were cultured for 48 hours in a minimal medium that contains glucose, mucin, or representative GAG (i.e., chondroitin sulfate C (CSC), HA, or HP) as a sole carbon source. Microbes in independent samples from three donors (one twenties, one thirties, and one fifties) has exhibited vigorous growth in each medium (*SI Appendix*, Fig. S3), indicating that microbes are present to degrade and/or assimilate GAG or mucin. The collected microbiota samples before and after growth were subjected to 16S rDNA amplicon sequence analysis (Fig. 1). Prior to cultivation, each sample had a wide variety of bacteria (Fig. 1*A*): Donors A, B, and C comprised of *Bifidobacterium, Prevotella*, and *Bacteroides* as the major genera, respectively. *Collinsella, Blautia, Faecalibacterium*, and *Dialister* genera were also detected well. The frequency of *Bacteroides* species ranged from 1.9 to 27.2%; among them, both *B. vulgatus* and *B. dorei* were most commonly observed. Succeeding cultivation for 24 hours in the presence of glucose, various genera such as *Bacteroides, Bifidobacterium, Erysipelatoclostridium*, and *Escherichia* propagated, even though many other early genera or species decayed. In a mucin-containing medium, *Collinsella* and *Blautia* genera also increased or maintained the frequency. Contrarily, only the *Bacteroides* species, especially *B. ovatus* and *B. xylanisolvens*, predominated the community (over 50% of total amplicons) in a GAG-containing medium in any donor sample. In view of the low initial frequency of both *B. ovatus* and *B. xylanisolvens* (0.0–1.7% and 0.0–0.6%, respectively), these species exhibited the best adaptive fitness to the used experimental conditions with GAG. Altogether, our data indicate that carbon sources are the key determinants in microbiota formation and maintenance.

**Fig. 1.**
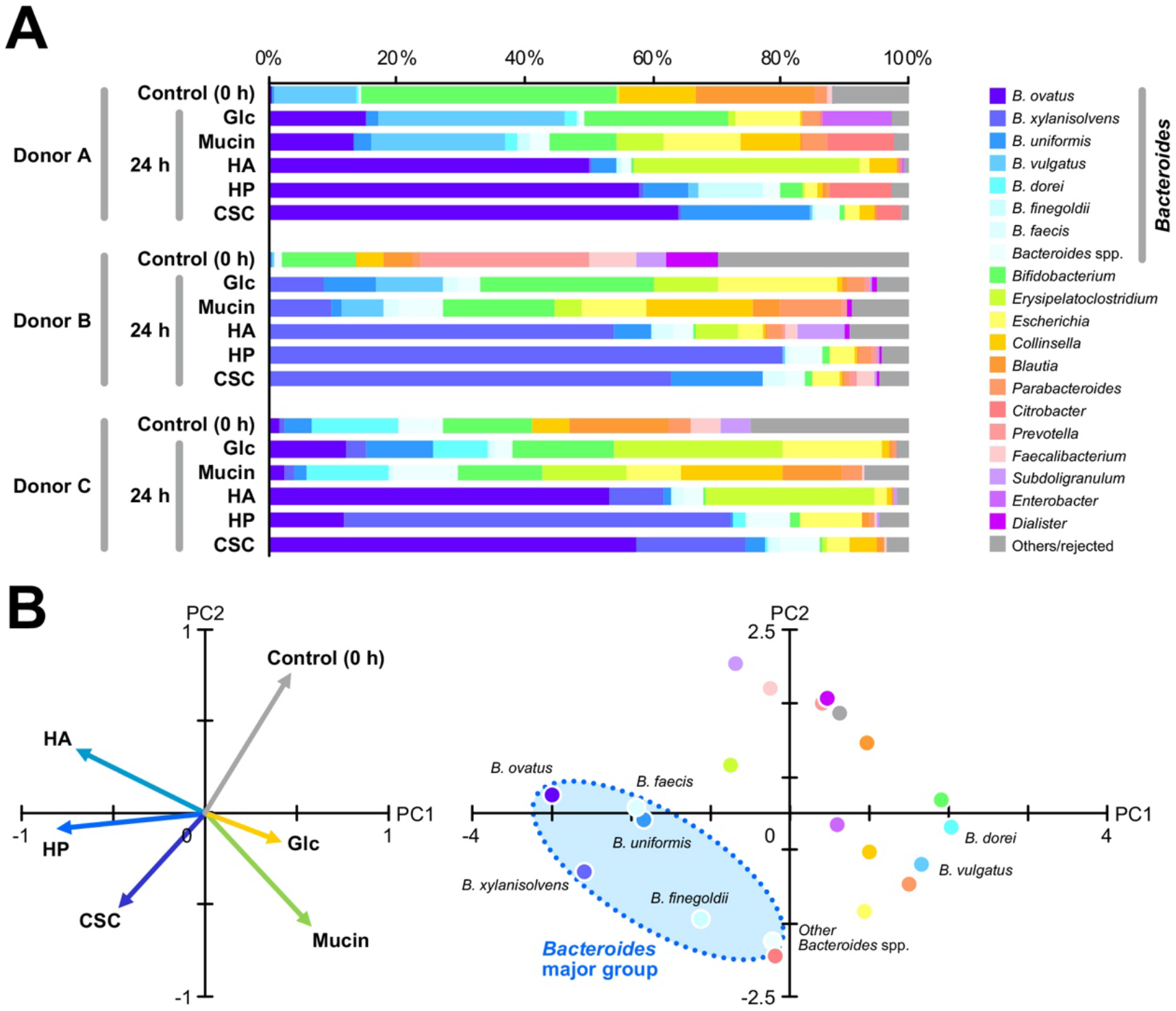
Human gut microbiota changes in the presence of mucosubstances. (*A*) Genera (or *Bacteroides* species) frequency profiles of the fecal samples cultivated in the presence of a sole carbon source (Glc, mucin, HA, HP, or CSC). (*B*) Principal component analysis of the microbiota profiles. The factor loading plot (*left*) and the PCA score plot (*right*) are seen. For marker colors in the PCA score plot, see the symbol legend in (*A*).

Principal component analysis (PCA) was done with the use of 0- and 24-hour data of averaged genus (or *Bacteroides* species) frequency (Fig. 1*B*). The first and second principal components (PC1 and PC2) accounted for 35.0% and 23.3% of the variance, respectively. Both the factor loading plot and the PCA score plot have shown that the majority of *Bacteroides* species, such as *B. ovatus, B. xylanisolvens, B. uniformis, B. finegoldii*, and *B. faecis*, formed a cluster with large negative PC1 values, corresponding to the high frequency in CSC, HA, or HP-containing medium. Therefore, GAG may be one of the reasons for the dominancy of most *Bacteroides* species in gut microbiota. The growth profiles of six major genera, exhibited from the growth curves of whole microbiota and the genus frequency in the microbiota, further supported that *Bacteroides* species prefer GAG than glucose or mucin (*SI Appendix*, Fig. S4), although, based on Fig. 1*B*, both *B. vulgatus* and *B. dorei* were plotted apart from the major *Bacteroides* group. The frequency of these two species was relatively high prior to cultivation but was lesser when GAG was used as a carbon source. Mucin was the only carbon source to grow the *Bacteroides* as well as five other major genera (*SI Appendix*, Fig. S4).

### Most *Bacteroides* Degrade GAGs

Since *Bacteroides* species prefer GAGs, their GAG-degrading abilities were assessed with the use of CSC, HA, and HP. We have earlier reported the degradation of GAGs by 6 *Bacteroides* species of 11 species tested through the halo assay with the use of GAG minimal medium plates (9). The detection of halo means there is a degradation of GAGs (27). Thus, to assess GAG degradation as one of the common characteristics among *Bacteroides* species, the other 17 species were also subjected to the halo assay (*SI Appendix*, Fig. S5), thus finding that *B*. *caccae, B. faecis*, and *B*. *finegoldii* degrade CSC. Additionally, *B*. *caccae, B*. *faecis, B. finegoldii, B*. *nordii*, and *B*. *xylanisolvens* were found to be novel HA-degrading *Bacteroides*. Contrarily, the species tested here do not degrade HP, showing that *Bacteroides* species prefer both CSC and HA containing 1,3-glycoside bond than HP containing 1,4-glycoside bond.

We then conducted the halo assay for the confirmation of the GAG-degrading ability of *Bacteroides* species under high nutrition conditions (Fig. 2). The nutrient-rich medium (Gifu Anaerobic Medium: GAM) plate was used for the halo assay since some *Bacteroides* species showed little growth in the GAG minimal plate, due to the lack of essential components including cysteine which is used for a reducing agent. In addition to 17 species used for the halo assay with GAG minimal plate, 8 other species were investigated with the use of the GAM plate. Because there was no precipitation formed in the HA-containing GAM plate, HA was not appropriate for this halo assay under high-nutrition conditions. In addition to CSC-degrading *Bacteroides* species found on the minimal medium plate, six species were found to degrade CSC: *B*. *cellulosilyticus, B*. *eggerthii, B*. *fluxus, B*. *gallinarum, B*. *massiliensis*, and *B*. *oleiciplenus*. However, in the case of HP minimal medium, there were no *Bacteroides* species found to degrade HP; the HP-containing GAM plate formed a halo, so that nine species exhibited HP-degrading ability as follows: *B*. *clarus, B. coprosuis, B*. *eggerthii, B*. *faecis, B*. *finegoldii, B*. *intestinalis, B*. *ovatus, B. stercoris*, and *B*. *thetaiotaomicron*. All in all, 18 of 28 *Bacteroides* species that were tested showed GAG (CSC, HA, and/or HP)-degrading ability (*SI Appendix*, Table S1).

**Fig. 2.**
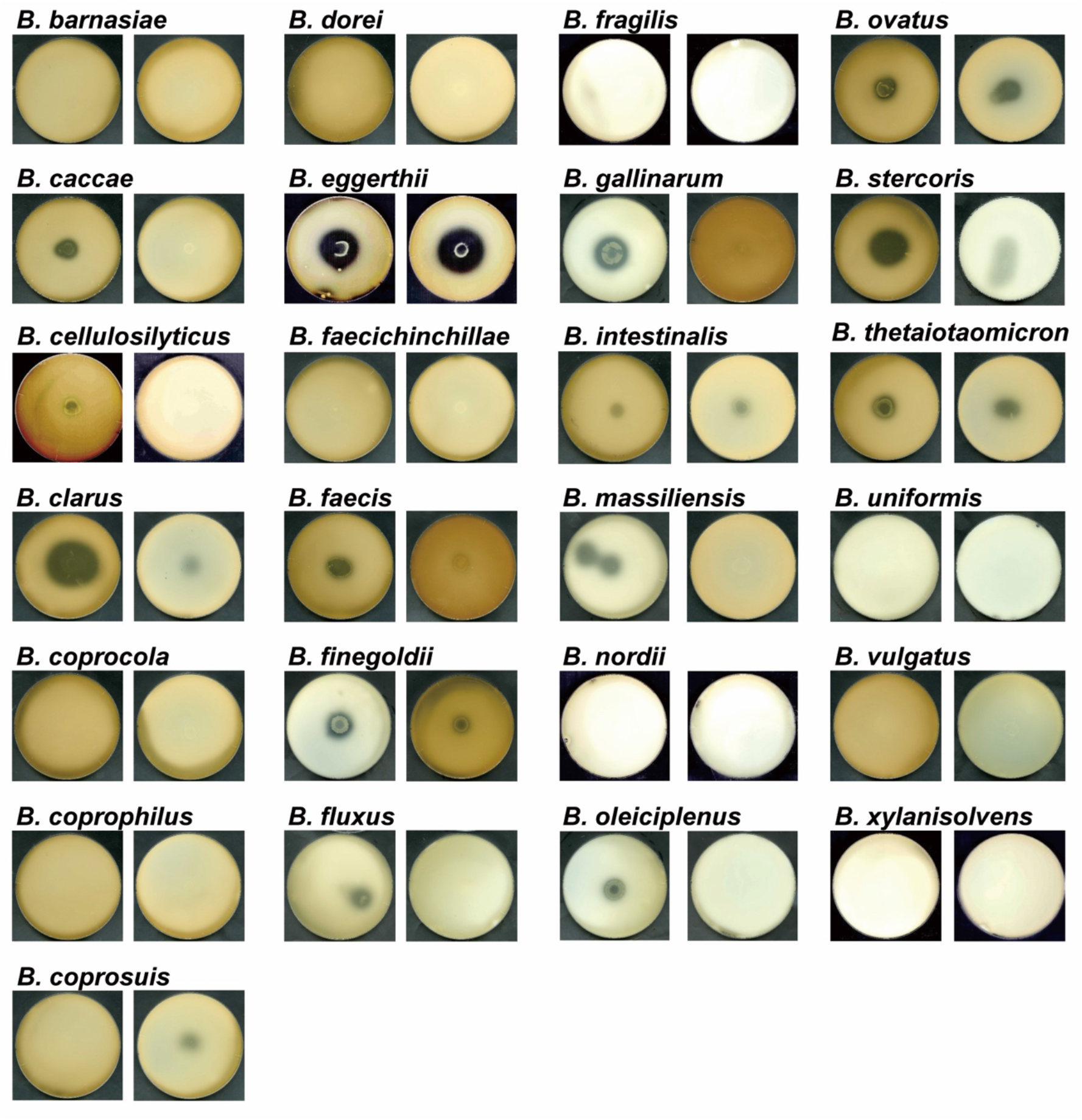
Degradation of GAGs by Bacteroides species. The halo assay for GAG degradation after 7-day incubation with *Bacteroides* species. CSC (*left*) or HP (*right*) was part of the nutrient-rich medium plate.

### Assimilation of GAGs by *Bacteroides* Species

The assimilation of GAGs by 29 *Bacteroides* species was further investigated by monitoring the optical density at 600 nm (OD_600_) of the bacterial culture in the liquid minimal medium such as CSC, HA, or HP as a sole carbon source (Fig. 3a, *SI Appendix*, Fig. S6), resulting in the growth of 20 species in the presence of CSC as follows: *B. barnesiae*, *B. caccae*, *B. cellulocsilyticus*, *B. clarus*, *B. coagulans*, *B. coprocola*, *B. coprophilus*, *B. eggerthii*, *B. faecis*, *B. finegoldii*, *B. fluxus*, *B. gallinarum*, *B. intestinalis*, *B. massiliensis*, *B. nordii*, *B. oleiciplenus*, *B. ovatus*, *B. stercoris*, *B. thetaiotaomicron*, and *B. xylanisolvens*. In the case of HA, 15 species proliferated, that is, *B. barnesiae*, *B. caccae*, *B. clarus*, *B. coagulans*, *B. coprocola*, *B. coprophilus*, *B. faecis*, *B. finegoldii, B. fluxus*, *B. gallinarum*, *B. massiliensis*, *B. nordii*, *B. ovatus*, *B. thetaiotaomicron*, and *B. xylanisolvens*. HP was assimilated by 13 species as follows: *B. clarus*, *B. eggerthii*, *B. faecis*, *B. finegoldii*, *B. fluxus*, *B. gallinarum*, *B. intestinalis*, *B. massiliensis*, *B. nordii*, *B. ovatus*, *B. stercoris*, *B. thetaiotaomicron*, and *B. xylanisolvens*. There is no species that has shown growth without GAGs, showing that these species grew by assimilating GAGs as a sole carbon source.

**Fig. 3.**
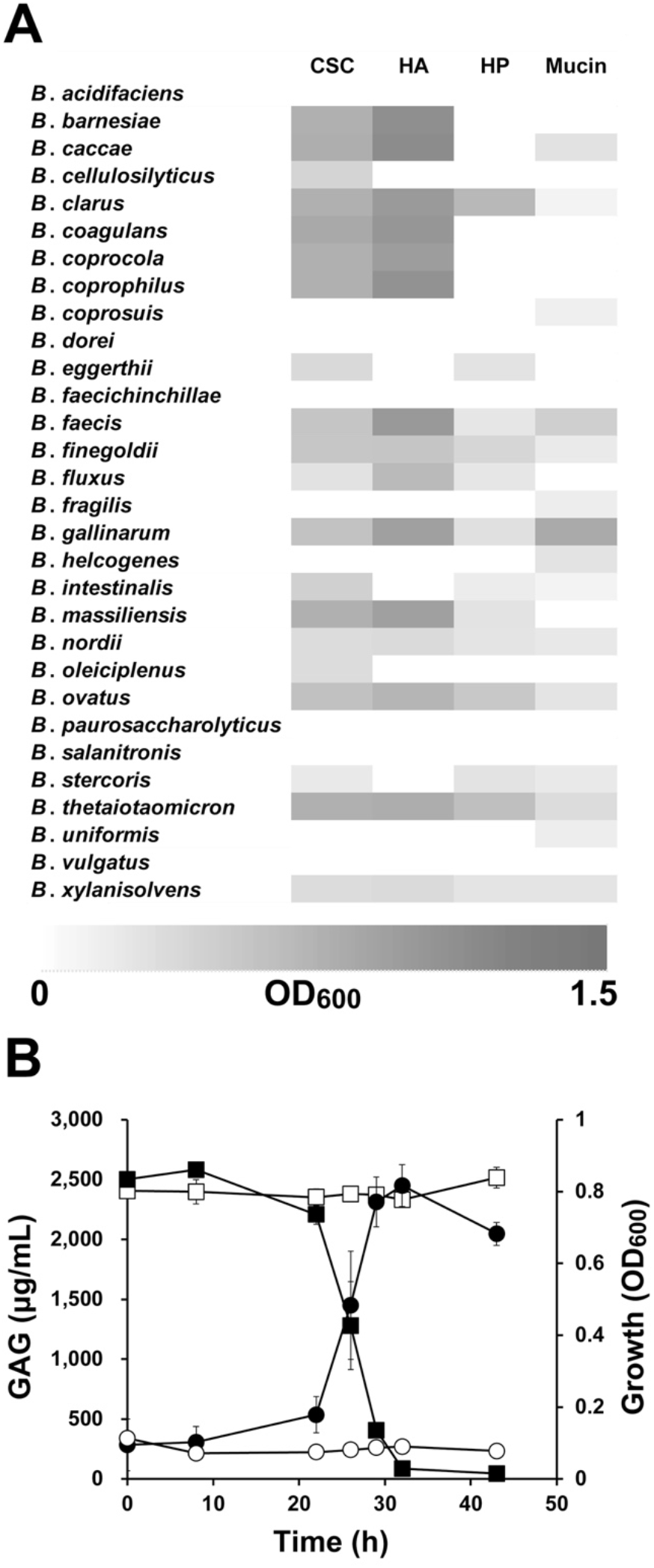
Assimilation of GAGs by *Bacteroides* species. (*A*) Growth level (OD_600_) of *Bacteroides* species in GAG (CSC, HA, or HP) or mucin minimal medium is shown using the heat map with OD_600_ from 0 (white) to 1.5 (black). (*B*) *Bacteroides faecis* growth-dependent degradation of CSC. Closed and open circles represent OD_600_ values of the culture broth in the presence and absence of *B. faecis*, respectively. Closed and open squares are the CSC concentrations of the culture broth in the presence and absence of *B. faecis*, respectively. Error bars show standard deviations (n = 3).

The time-dependent decrease of GAG concentration in the CSC minimal medium was examined during the growth of *B. faecis* as a representative. In accordance to the OD_600_ increase of the bacterial culture broth, there was a concentration decrease of CSC, and the polysaccharide was finally totally consumed (Fig. 3*B*). This result has directly shown that CSC was assimilated by *B. faecis*. In total, 20 species of 29 *Bacteroides* species that were tested here assimilate GAGs (*SI Appendix*, Table S1).

In the majority of *Bacteroides* species, the growth profiles in human gut microbiota (Fig. 1*B*) and in pure culture (Fig. 3*A*) were quite steady: *B. ovatus, B. xylanisolvens, B. finegoldii*, and *B. faecis* in the major *Bacteroides* group showed high assimilation ability, while *B. vulgatus* and *B. dorei* did not have GAG utilization. However, a major-group species (*B. uniformis*) has no degradation or assimilation of GAG. Such species could not feed by themselves on GAG but may grow in microbiota with the help of GAG-degrading/ GAG-assimilating bacteria.

### Frequent Detection of GAG Lyase Gene in Human Gut Microbiota

As a first step of bacterial action on GAGs, bacterial GAG lyases are very important for depolymerization of GAGs (*SI Appendix*, Fig. S1). Among these, HA/CS and heparan sulfate lyases are categorized as Polysaccharide Lyase Families 8 (PL8) and PL12, respectively, in the database CAZy (http://www.cazy.org/) (6, 16, 17). The primers to amplify these lyase genes were designed by referring to those genes in four *Bacteroides* species as follows: *B. cellulosilyticus*, *B. ovatus*, *B. thetaiotaomicron*, and *B. xylanisolvens*. Family PL8 and PL12 GAG lyase genes were discovered from three independent feces, and their frequency was 6.0%, 0.85%, and 3.2% in human gut microbiota on the basis of the bacterial cells calculated from 16S rDNA amplification (*SI Appendix*, Table S2). These values were reasonable based on the frequency of *Bacteroides* species in fecal samples as described above.

### Assimilation of Mucin by *Bacteroides* Species

Besides GAG, the assimilation of mucin was assessed with the use of 28 *Bacteroides* species on the purified mucin minimal medium. Although no *Bacteroides* species showed a significant growth on the medium without mucin, 15 species assimilated mucin as follows: *B. caccae*, *B. clarus*, *B. coprosuis*, *B. faecis*, *B. finegoldii*, *B. fragilis*, *B. gallinarum*, *B. helcogenes*, *B. intestinalis*, *B. nordii*, *B. ovatus*, *B. stercoris*, *B. thetaiotaomicron*, *B. uniformis*, and *B. xylanisolvens* (Fig. 3*A*, *SI Appendix*, Table S1). Among the mentioned species, six, *B. clarus*, *B. coprosuis*, *B. faecis*, *B. gallinarum*, *B. helcogenes*, and *B. nordii*, were the first ones to show mucin assimilation. Since commercially available mucin contained GAGs, the amount of sulfated GAGs in the purified mucin was measured using the DMMB method, resulting in the purified mucin containing 7–8% (w/v) of GAGs, which corresponds to 0.02% (w/v) GAGs included in 0.25% (w/v) mucin medium which is used for the growth assay. To remove the possibility that *Bacteroides* species assimilated only GAG in the mucin minimal medium, *B. thetaiotaomicron* was cultured in 0.25% (w/v) mucin or 0.02% (w/v) GAG minimal medium. As a consequence, there was a higher growth level on mucin than that on GAG (*SI Appendix*, Fig. S2), showing that the bacterial cells assimilated mucin.

### *Bacteroides* Species Utilize GAG or Mucin as a Sole Nitrogen Source

Nitrogen is one of the essential elements, despite its limited quantity in the gut environment. Therefore, bacteria need to acquire nitrogen to be indigenous in the colon. Because nitrogen is included in GAG or mucin, we tested whether *Bacteroides* species can use HA or mucin as a sole nitrogen source (*SI Appendix*, Fig. S7). Three species, *B. thetaiotaomicron, B. ovatus*, and *B. faecis*, have shown a remarkable growth even in nitrogen-restricted GAG or mucin medium. In the meantime, these three species showed no growth in nitrogen-restricted glucose medium, even though they grew well in the presence of ammonium sulfate. *Bacteroides* species can, therefore, use GAG or mucin as carbon and nitrogen sources. This feature is one of the reasons why *Bacteroides* species have dominance in the gut environment.

### Metabolites from *Bacteroides* Species during Assimilation of Host Mucosubstances

Metabolites in the culture broth secreted by *B. thetaiotaomicron* or *B. ovatus* were investigated in the GAG (HA) or mucin minimal medium, resulting in the detection of 19 kinds of standard amino acids, γ-amino butyrate (GABA), α-amino butyrate, and ethanol amine (Fig. 4*A*). A huge amount of Ala was seen under all conditions, and GABA, Glu, and Val were remarkably secreted when HA was assimilated. On the contrary, when the bacterial cells assimilated mucin, there was an excretion of basic amino acids, for example, Lys and Arg. Moreover, organic acids were also investigated. Three *Bacteroides* species, *B. thetaiotaomicron*, *B. ovatus*, and *B. faecis*, grown in the HA minimal medium secrete short-chain fatty acids including acetic and propionic acids as well as lactic, succinic, and/or formic acids (Fig. 4*B*).

**Fig. 4.**
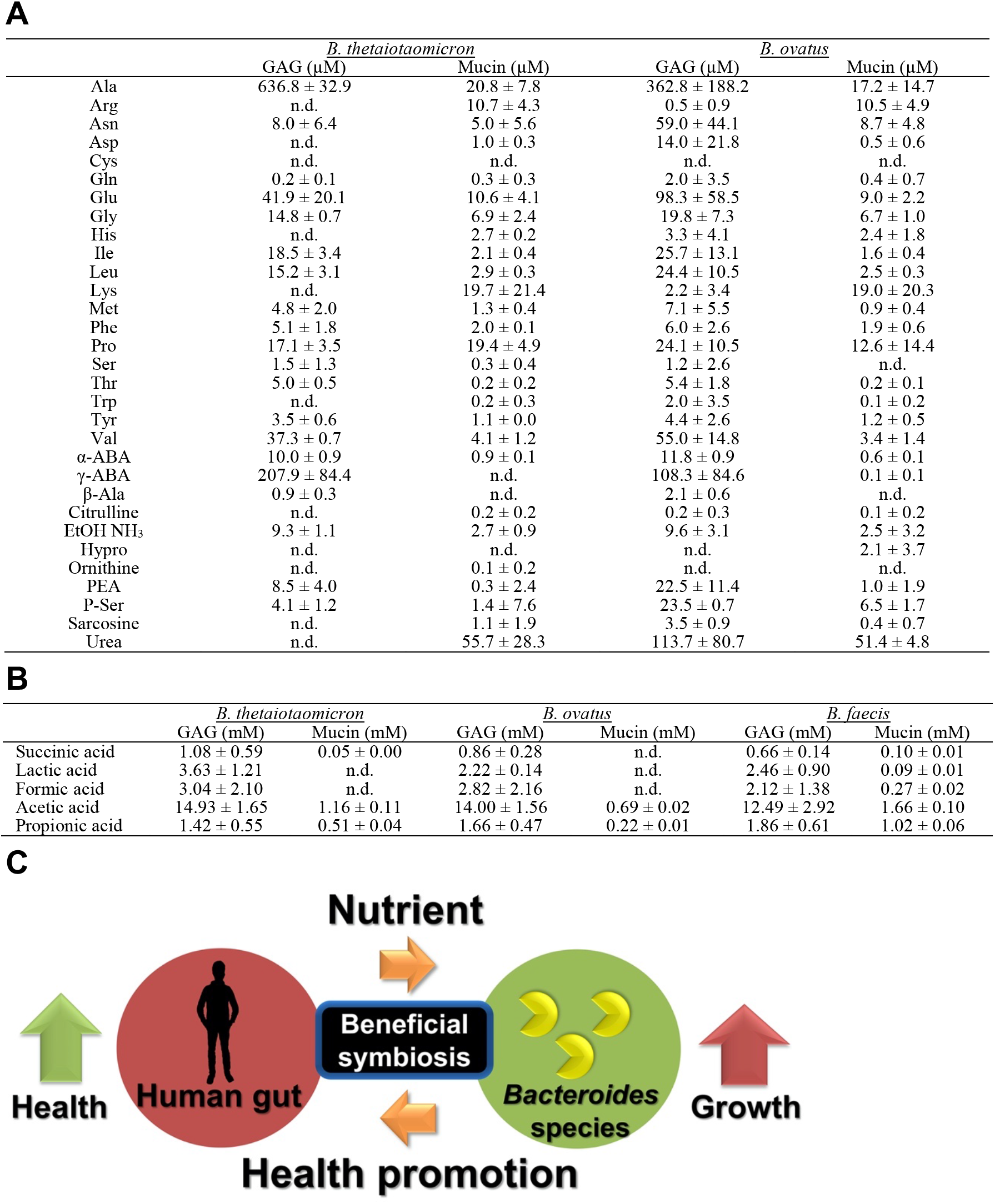
Mutually beneficial symbiosis between human and *Bacteroides* species. (*A*) Secretion of amino acids from GAG (HA) and mucin by *Bacteroides* species. n.d., not detected. (*B*) Secretion of organic acids from GAG (HA) and mucin by *Bacteroides* species. n.d., not detected. (*C*) Symbiosis model. The human provides *Bacteroides* species in the gut with extracellular mucosubstances (GAGs and mucin) for their nutrition and residence place. *Bacteroides* species dominate gut microbiota by assimilating host mucosubstances as a carbon/nitrogen source and secrete molecules including amino acids and short-chain fatty acids which are essential in human health promotion.

## Discussion

Based on the above-described results, we have discussed the following: Since GAGs and mucin are constantly supplied in human gut independent from nutrient intake by the host, GAG or mucin assimilation abilities are necessary for survival of dominant bacteria in the human gut, supporting the hypothesis that assimilation of GAG or mucin is essential for gut microbiota dominance in the human gut. In fact, the abundance of *Bacteroides* species in the gut microbiota has increased in female mice by oral administration of chondroitin sulfates A and C (28). The gene expression involved in the assimilation of GAGs is also upregulated in *Bacteroides* species in the presence of GAGs (18, 29). As regards structure, GAG is simpler than mucin, so that GAG may be degraded easier than mucin (3, 30). Indeed, more species have shown an assimilation ability toward GAG rather than mucin (Fig. 3*A*, *SI Appendix*, Table S1).

Amino acids secreted by gut microbiota are used by the host as well as other gut bacteria (31). *Bacteroides* species grown in GAG or mucin minimal medium secreted most of the standard amino acids and organic acids such as short-chain fatty acids (Fig. 4), showing that the host human and other bacteria can assimilate these metabolites. Additionally, *Bacteroides* species could provide important amino acids to host human by converting host GAGs to those amino acids which are not dependent from nutrient intake by the host. Even with the weight of gut microbiota of about 1.5 kg and *Bacteroides* species being dominant species (~50% of total), the amount of secreted amino acids is not negligible, even though amino acids are generally absorbed by the small intestine and *Bacteroides* species are broadly distributed from the stomach to the large intestine (32). The quantities of amino acids produced by *Bacteroides* species are almost equal to the recommended amount for a person on a day by the WHO (*SI Appendix*, Table S3), indicating that the human obtains a considerable amount of nutrition from metabolites secreted by gut microbiota. In the recent time, *Bacteroides* species produce much GABA, and negative correlation is apparent between their abundance and depressive symptoms (33), showing that GABA which is produced by human gut microbiota is as a signal molecule in the enteric nervous system (34). Thus, besides nutritional effects, *Bacteroides* species producing GABA from the host GAG may be a contributing factor to human mental health.

Furthermore, gut microbiota is involved in host immunity and/or energy source via the secretion of organic acids such as short-chain fatty acids (19). For the most part, short-chain fatty acids which are very important to the host are produced at 50–100 mM in the colonic lumen by gut microbiota (35, 36). Based on the secretion level from the mucosubstances (GAG and mucin) (Fig. 4*B*), *Bacteroides* species are the main producers of short-chain fatty acids in the human gut. Since the intestine mucus of the host is supplied inside of the gut which is independent from the nutrient uptake by the host, *Bacteroides* species should supply physiologically essential molecules to the host or other bacteria under fasting conditions. These results support the fact that *Bacteroides* species are known as the next-generation probiotics (15, 22) and/or pharmabiotics (34).

The overall results acquired here hypothesized that there is a mutually beneficial relationship model between *Bacteroides* species and the human host (Fig. 4*C*). It is essential for humans to secrete extracellular gut mucosubstances including GAGs and mucin for the maintenance of cell structure/function and prevention of inflammation due to pathogenic bacteria. Some gut bacteria convert these mucosubstances to essential molecules for the host. This mutual beneficial relationship has probably been accepted as a result of natural selection in the human gut. For humans, *Bacteroides* species in the gut leads to the promotion of host health by providing amino acids and organic acids such as short-chain fatty acids, so that the human immune system has not excluded *Bacteroides* species from the gut. On the contrary, for the *Bacteroides* species, the human gut is a well-organized place to survive due to the continuous supply of large quantities of mucosubstances in which *Bacteroides* species can assimilate dominantly in human gut microbiota. Human health is also necessary for *Bacteroides* species to keep the residence place safe.

Finally, the mutually beneficial symbiosis between humans and gut-dominant *Bacteroides* species is posited via bacterial assimilation of host mucosubstances and secretion of metabolites for host health promotion.

## Materials and Methods

### Materials

HA and DMMB were purchased from Sigma-Aldrich. Mucin from porcine stomach and CSC were acquired from Wako. HP was from Nacalai Tesque. The nutrition-rich medium, GAM, was from Nissui Pharmaceutical Co. All other analytical grade chemicals utilized in this study were available commercially. The feces were kindly given to us as gifts by the Japanese volunteers. Informed consent was acquired from all subjects, and experiments with the use of these feces were approved by the Committee of Research Activity Promotion of Graduate School of Agriculture, Kyoto University.

### Purification of GAG and Mucin

Low-molecular-weight molecules contaminated in each GAG reagent were removed by dialysis against pure water. Unless otherwise stated, the percent concentration represents w/v. Mucin was dissolved at a 2% final concentration in 20 mM potassium phosphate buffer (pH 7.8), which contains 0.1 M NaCl. Several drops of toluene were placed in the mucin suspension, which is followed by stirring for 1 hour at room temperature. The suspension was then adjusted to a pH of 7.2 with 2 M NaOH and stirred for 23 hours at room temperature. The supernatant was collected after centrifugation (4°C, 10,000×g, 10 min). Cold ethanol was then added to become 60% (v/v), for the precipitation of mucin to occur. Centrifugation was done with the same condition, and the resultant pellet was then dissolved in 0.1 M NaCl. The addition of ethanol followed by centrifugation was repeated twice. The last step was to dissolve the pellet in pure water (80 mL) instead of 0.1 M NaCl. This solution was dialyzed against pure water and then freeze-dried to obtain the purified mucin (37).

### Bacteria Strains

A total of 30 *Bacteroides* species was utilized in this study. Among them, 29 species were purchased from Japan Collection of Microorganisms (JCM), such as *B. acidifaciens* JCM10556, *B. barnesiae* JCM13652, *B. caccae* JCM9498, *B. cellulosilyticus* JCM15632, *B. clarus* JCM16067, *B. coagulans* JCM12528, *B. coprocola* JCM12979, *B. coprophilus* JCM13818, *B. coprosuis* JCM13475, *B. dorei* JCM13471, *B. eggerthii* JCM12986, *B. faechinchillae* JCM17102, *B. faecis* JCM16478, *B. finegoldii* JCM13345, *B. fluxus* JCM16101, *B. fragilis* JCM11019, *B. gallinarum* JCM 13658, *B. helcogenes* JCM6297, *B. intestinalis* JCM13265, *B. massiliensis* JCM13223, *B. nordii* JCM12987, *B. oleiciplenus* JCM16102, *B. ovatus* JCM5824, *B. paurosaccharolyticus* JCM15092, *B. salanitronis* JCM13657, *B. stercoris* JCM9496, *B. thetaiotaomicron* JCM5827, *B. uniformis* JCM5828, and *B. xylanisolvens* JCM15633.

In addition to this*, B. vulgatus* NBRC14291 was from the Biological Resource Center in the National Institute of Technology and Evaluation (NBRC/NITE).

### 16S rDNA Amplicon Sequence Analysis

There is approximately 1 g of human fecal sample dissolved in 10 mL of sterilized saline (0.9% NaCl). A 300-μL aliquot of this suspension was inoculated into a test tube which contains 15 mL of assimilation validation liquid medium [0.1% ammonium sulfate, 0.226% KH_2_PO_4_, 0.09% KH_2_PO_4_, 0.0004% FeSO4(II), 0.09% NaCl, 0.0027% CaCl_2_/2H_2_O, 0.002% MgCl_2_/6H_2_O, 0.001% MnCl_2_/4H_2_O, 0.001% CoCl_2_, 0.0005% hemin, 0.00001% vitamin K1, 0.00001% ethanol, 0.0000005% vitamin B12, and 0.04% L-cysteine, with or without 0.25% glucose, dialyzed GAG (CSC, HA, or HP) or the purified mucin]. After being anaerobically cultured at 37°C for 24 or 48 hours, cells acquired after centrifugation were washed using the sterilized saline and immediately frozen in liquid nitrogen.

Both DNA extraction from human gut microbiota and 16S rDNA amplicon sequence analysis were performed by TechnoSuruga Laboratory Co., based on a previously reported method (38). In summary, the V3–V4 region of 16S rDNA was amplified with the use of the 341F/R806 primer sets. Sequencing was conducted using a paired-end, 2×300-bp cycle run on a MiSeq sequencing system (Illumina) and MiSeq Reagent Kit version 3 (600 cycle) chemistry. After the sequencing was done, image analysis, base calling, and error estimation were done with the use of the Illumina Real-Time Analysis software (version 1.17.28). Paired-end sequencing with read lengths of approximately 430 bp was done as well. Succeeding the demultiplexing, a clear overlap in the paired-end reads was seen. This made paired reads be joined together with the fastq-join program. Only reads that had quality value (QV) scores of ≥ 20 for more than 99% of the sequence were extracted for supplemental analysis. Metagenome@KIN software (World Fusion) was utilized for homology searching with the determined 16S rDNA sequences, against the DB-BA13.0 microbial identification database (TachnoSuruga Laboratory) (39, 40). Bacterial species were then identified based on the data from 97% similarity cut-off with DB-BA13.0. All sequences have been deposited in the DNA Data Bank of Japan (DDBJ) under the accession number DRA010273.

### GAG Quantitation

Fecal samples (about 2 g) acquired from three volunteers were dissolved in 50 mL of pure water followed by overnight rotation. Centrifugation and filtration were done for insoluble residues to be removed. The reagent DMMB determined the GAG content at a final concentration of 0.0016% in 0.304% glycine, 0.16% NaCl, and 0.057% (v/v) acetic acid as quantitative reagent (26). Standard curve was made using a variety of CSC concentrations (0, 10, 20, 30, 40, and 50 mg/mL). The absorbance at 525 nm (Abs_525_) was measured after 80 μL of samples and 800 μL of quantitative solution were mixed. To remove the effect of other contaminated materials toward Abs_525_ in feces, Abs_525_ values of tenfold diluted fecal samples with pure water were measured and subtracted from Abs_525_ values of DMMB when GAGs in feces were quantified.

### Halo Assay for the Detection of GAG Degradation

*Bacteroides* species were grown in an anaerobic condition at 37°C overnight in 5 ml of liquid GAM (1% peptone, 0.3% soy peptone, 1% proteose peptone, 1.35% digested serum powder, 0.5% yeast extract, 0.22% meat extract, 0.12% liver extract, 0.3% glucose, 0.25% KH_2_PO_4_, 0.3% NaCl, 0.5% soluble starch, 0.03% L-cysteine hydrochloride, and 0.03% sodium thioglycolate) as a preculture. The preculture broth (1 mL) was centrifuged at 9,700×g for 5 min, and the resulting pellet (bacterial cells) was then washed with 1 mL of sterilized saline (0.9% NaCl), which is followed by centrifugation (9,700×g, 5 min). The pellet was suspended in saline to acquire OD_600_ = 1. The cell suspension (10 μL) was seen on the center of the halo minimal medium plate [0.1% yeast extract, 0.1% KH_2_PO_4_, 0.1% Na_2_HPO_4_, 0.01% MgSO_4_/7H_2_O, 1% bovine serum albumin (BSA), 0.2% dialyzed GAG (CSC, HA, or HP), and 1% agar]. The plate was incubated in an anaerobic condition at 37°C for 7 days. Afterward, 1 mL of 2 M acetic acid was spread on the plate to form white precipitation of the BSA-GAG complex. In the case of GAG-degrading bacteria, halo (clear zone) was seen on the plate due to the lack of GAG polymer (27).

For some *Bacteroides* species with a little growth in the halo minimal medium plate, the nutrition-rich halo medium plate was also utilized. This plate has GAM to supply a proper reducing agent for anaerobic bacteria as follows: 0.4% peptone, 0.12% soy peptone, 0.4% proteose peptone, 0.54% digested serum powder, 0.2% yeast extract, 0.088% meat extract, 0.048% liver extract, 0.12% glucose, 0.1% KH_2_PO_4_, 0.12% NaCl, 0.2% soluble starch, 0.012% L-cysteine hydrochloride, 0.012% sodium thioglycolate, 1% BSA, and 0.2% dialyzed GAG (CSC or HP).

### Assimilation Assay

To investigate the mucosubstance assimilation by *Bacteroides* species, OD_600_ of the culture broth was measured after their inoculation into a variety of liquid media. *Bacteroides* species precultured in liquid GAM was centrifuged at 15,000×g for 10 min, and the resulting pellet (bacterial cells) was washed using sterilized saline (0.9% NaCl). The bacterial cells were washed thrice and finally suspended in saline to acquire OD_600_ = 1. The cell suspension (300 μL) was inoculated into 15 mL of the modified assimilation validation liquid medium (41) [0.1% ammonium sulfate, 0.226% KH_2_PO_4_, 0.09% KH_2_PO_4_, 0.0004% FeSO_4_ (II), 0.09% sodium chloride, 0.0027% CaCl_2_/2H_2_O, 0.002% MgCl_2_/6H_2_O, 0.001% MnCl_2_MH_2_O, 0.001% CoCl_2_, 0.0005% hemin, 0.00001% vitamin K1, 0.00001% ethanol, 0.0000005% vitaminB12, 0.04% L-cysteine, and 0.25% dialyzed GAG (CSC, HA, or HP) or the purified mucin] and was further cultured at a temperature of 37°C with anaerobic conditions. The negative control, validation liquid medium with no inoculation, was also incubated to check the background of OD_600_. For *B. faecis*, OD_600_ of the bacterial culture with CSC as a sole carbon source was monitored by measuring the CSC concentration.

Three species, *B. faecis*, *B. ovatus*, and *B. thetaiotaomicron*, were grown in nitrogen-restricted medium (assimilation validation liquid medium without ammonium sulfate) with HA, mucin, or glucose present as a sole carbon source to check whether these bacterial cells can use GAG or mucin as a sole nitrogen source. As indicated above, bacterial growth was monitored by measuring OD_600_ of the culture broth.

### Metagenomics of Gut Bacteria for GAG Degradation

Metagenomics were accomplished with the use of fecal samples from three volunteers. The analyses were conducted by TechnoSuruga Laboratory Co. DNA primers, for the amplification of GAG lyase genes, were designed by referring to family PL8 and PL12 lyase gene sequences found in database CAZy (http://www.cazy.org/) as follows: PL8 forward,

CTSGAYGGDGCMACVAAYATAGA; PL8 reverse,

TTTCCATCGGGAGWDCCRGCHAD; PL12 forward,

RAYTAYCCVGGWYTRGARAAAG; and PL12 reverse,

WCCAYTKATGRCGATGMADYTG. The total bacterial cell numbers were estimated by real-time PCR with the use of primers which are specific to 16S rDNA.

### Metabolic Assay of *Bacteroides* Species through Assimilation of GAG or Mucin

Three species, *B. faecis*, *B. ovatus*, and *B. thetaiotaomicron*, were cultured in the assimilation validation liquid medium that contained HA or mucin as a sole carbon source. Organic acids in the bacterial culture broth for 48 hours were measured by TechnoSuruga Laboratory Co. at 45°C of the column oven temperature and 0.8 mL/min of flow rate with the use of 5 mM *p*-toluenesulfonic acid as a solvent by HPLC system (Shimadzu) that is equipped with a column of Shim-pack SCR-102(H) and electrical conductivity detector of CDD-10A. As regards *B. thetaiotaomicron* and *B. ovatus*, the bacterial culture broth was also subjected to Amino Acid Analyzer L-8900 (Hitachi High-Technologies) which is equipped with a column of #2622SC-PF and a detector of photometer in the Global Facility Center of Hokkaido University. Free amino acids were seen at wavelengths of 570 and 440 nm by the post-staining method with ninhydrin. Before analyses, mucin was taken out with the use of ethanol precipitation. The medium with no cells was also investigated as a background.

## Acknowledgements

This work was supported in part by Grants-in-Aid for Scientific Research from the Japan Society for the Promotion of Science (to W.H.), and Research Grant (to W.H.) from Yakult Bio-science Foundation. The authors would like to thank Enago (www.enago.com) for the English language review.

## Author contributions

W.H. designed the study; M.S., K.K., T.K., R.T., and W.H. performed the experiments; M.S., K.K., T.K., D.W., R.T., and W.H. analyzed the data; M.S., D.W., R.T., and W.H. wrote the manuscript.

