## Supplementary Information for "Mutually Beneficial Symbiosis Between Human and Gut-Dominant *Bacteroides* Species Through Bacterial Assimilation of Host Mucosubstances"

**This PDF file includes:**

Figures S1 to S7

Tables S1 to S3

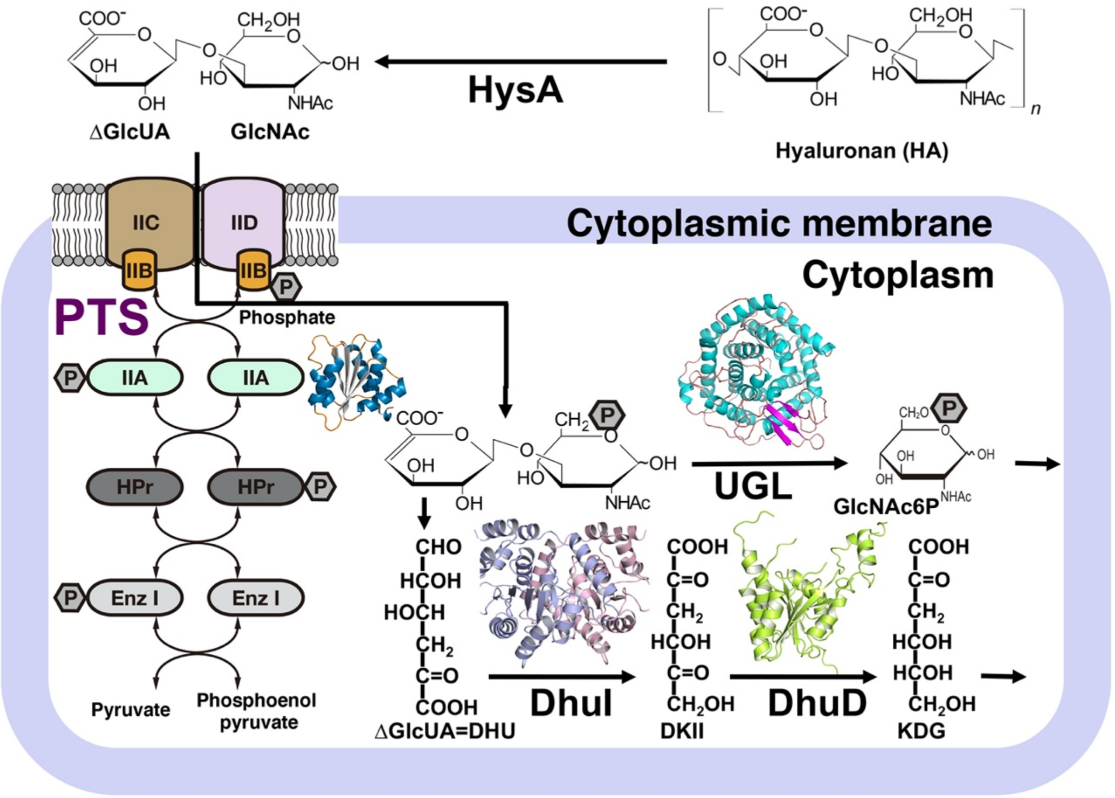


**Figure S1. Depolymerization, import, degradation, and metabolism of GAG by *Streptococcus agalactiae*.** Hyaluronan is depolymerized to unsaturated disaccharide (ΔGlcUA-GlcNAc) via the β-elimination reaction by extracellular hyaluronate lyase (HysA). The resultant disaccharide is imported to the cytoplasm via phosphorylation by the phosphotransferase system (PTS) and degraded to constituent monosaccharides (ΔGlcUA and GlcNAc-6-phosphate (GlcNAc6P)) by the unsaturated glucuronyl hydrolase (UGL). 4-Deoxy-L-*threo*-5-hexosulose-uronate (Dhu) nonenzymatically generated from ΔGlcUA is converted to 2-keto-3-deoxy-D-gluconate (KDG) via 3-deoxy-D-*glycero*-2,5-hexodiulosonate (DKII) through successive reactions of isomerase (DhuI) and NADH-dependent reductase/dehydrogenase (DhuD).

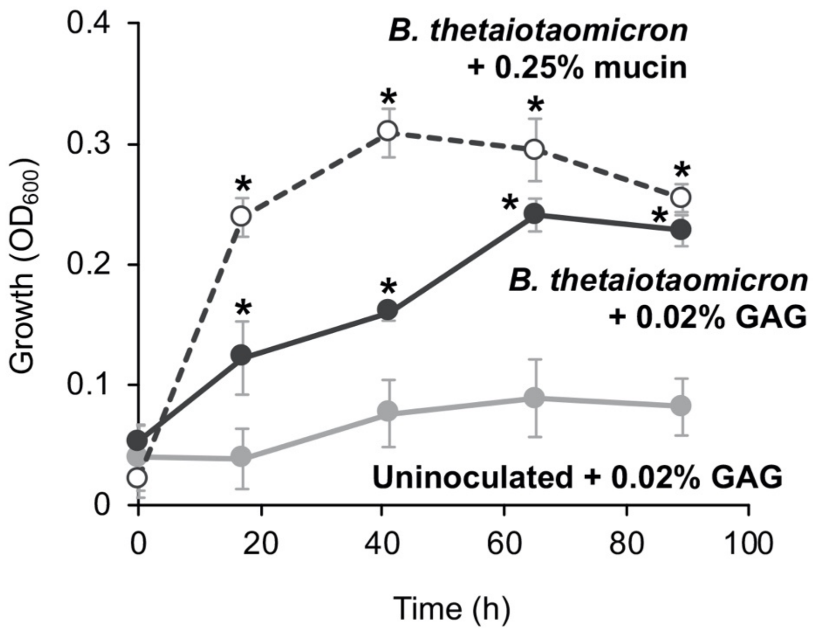


**Figure S2. Growth profile of *B. thetaiotaomicron* in 0.02% HA minimal medium or in 0.25% mucin minimal medium.** Each data point represents the mean and standard deviations from three independent experiments. Asterisks indicate significant differences from data of the bacterial cell-free medium ($p < 0.05$; *t*-test).

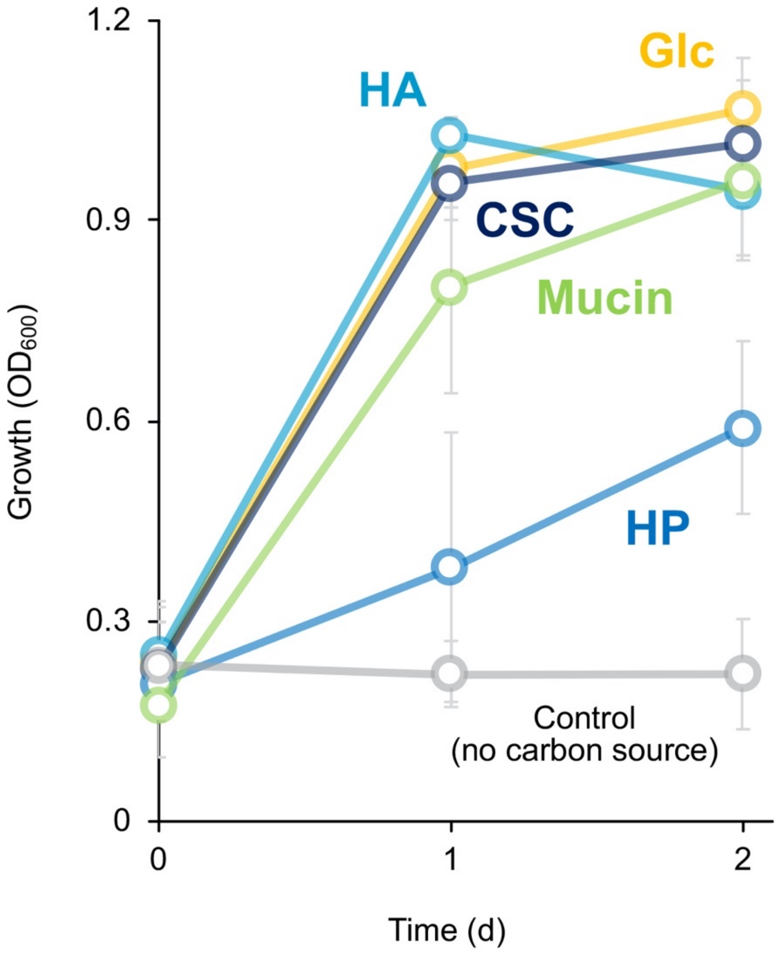


**Figure S3. Growth profile of human gut microbiota cultivated in the presence of a single carbon source (CSC, Glc, HA, HP, or mucin).** Data represent means and standard deviations from three independent experiments with various donors.

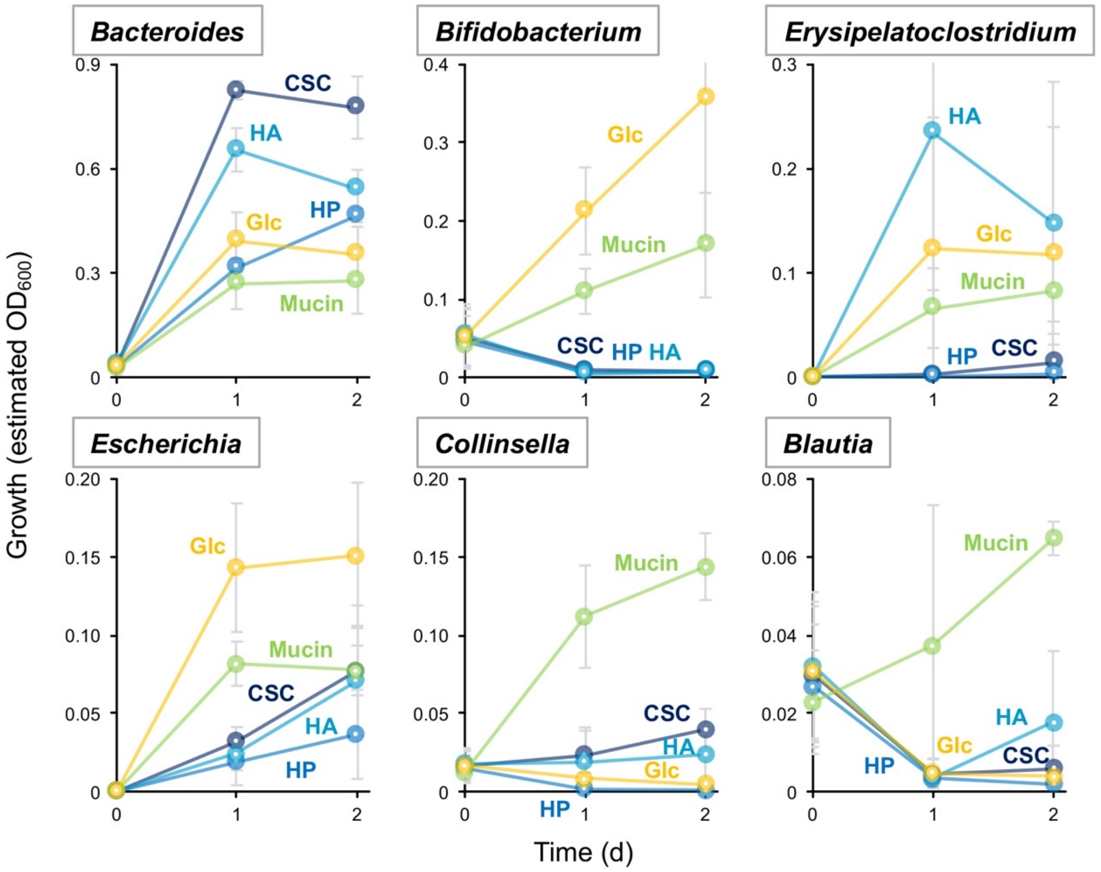


**Figure S4. Estimated growth profile of individual genera in human gut microbiota cultivated in the presence of a single carbon source.** Estimated $OD_{600}$ was calculated by multiplying $OD_{600}$ by each genus frequency in the microbiota. Data represent means as well as standard deviations from three independent experiments with various donors.

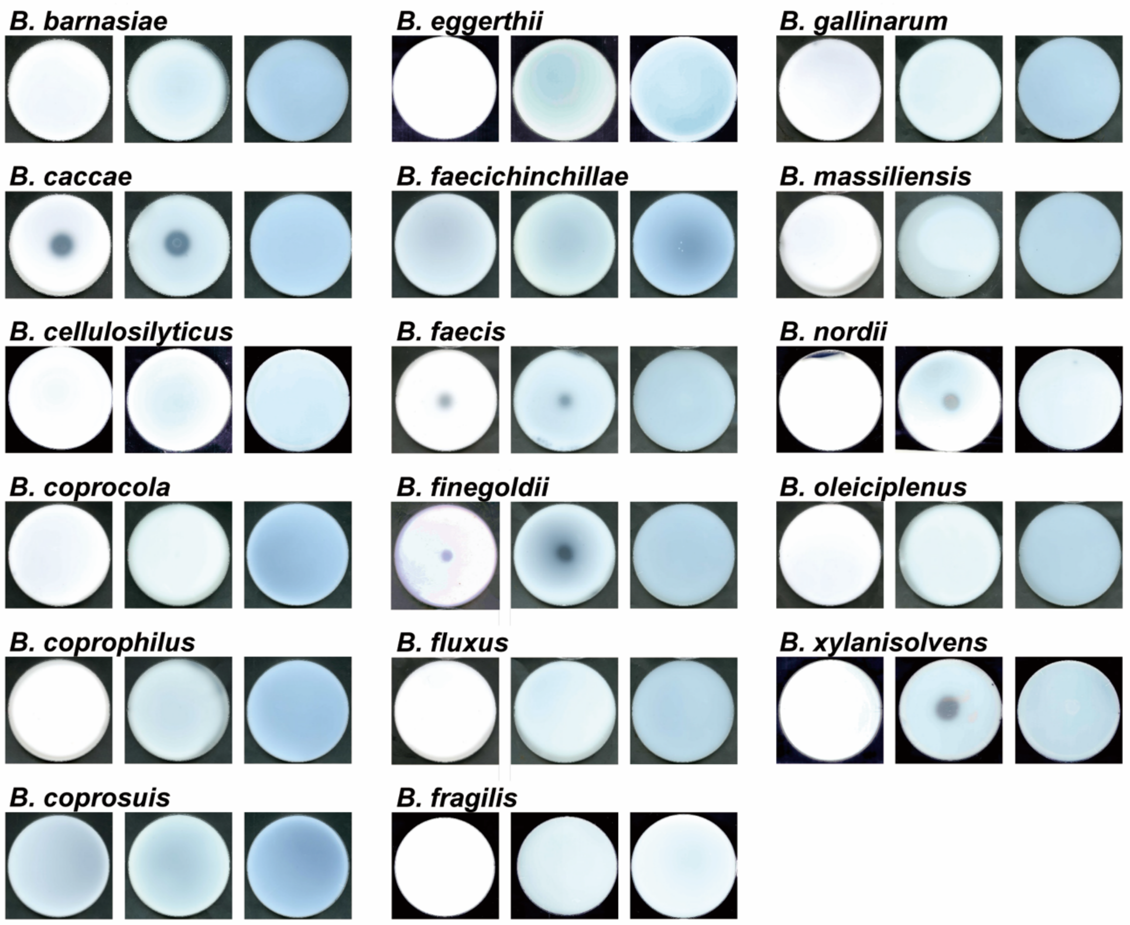


**Figure S5. Degradation of GAGs by *Bacteroides* species.** The halo assay for GAG degradation after incubation for 7 days with *Bacteroides* species. CSC (left), HA (center), or HP (right) was included in the minimal medium plate.

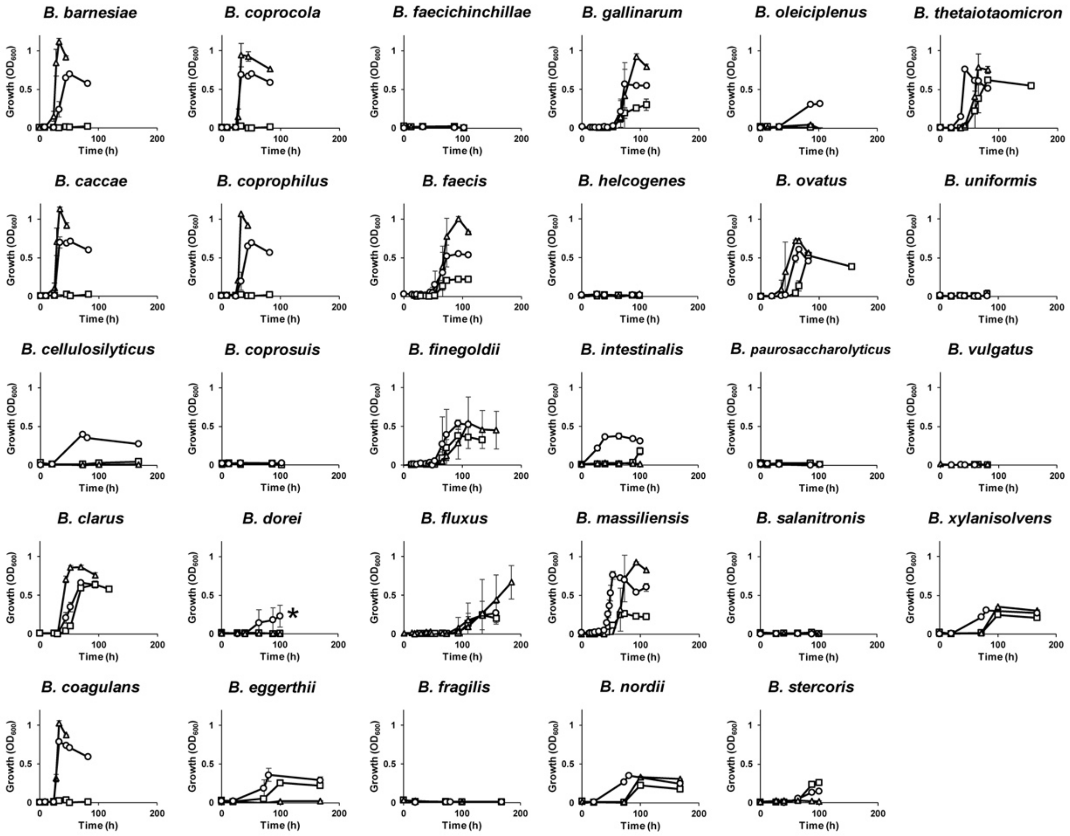


**Figure S6. Growth profiles of *Bacteroides* species in GAG minimal medium.** Each of the 29 *Bacteroides* species was inoculated in GAG (CSC, HA, or HP) minimal medium and $OD_{600}$ of the culture broth (CSC, circle; HA, triangle; HP, square) was periodically measured. Data represent means and standard deviations from three independent experiments. *The growth of *B. dorei* in the presence of CSC is not necessarily trustworthy because of large errors compared to $OD_{600}$ values.

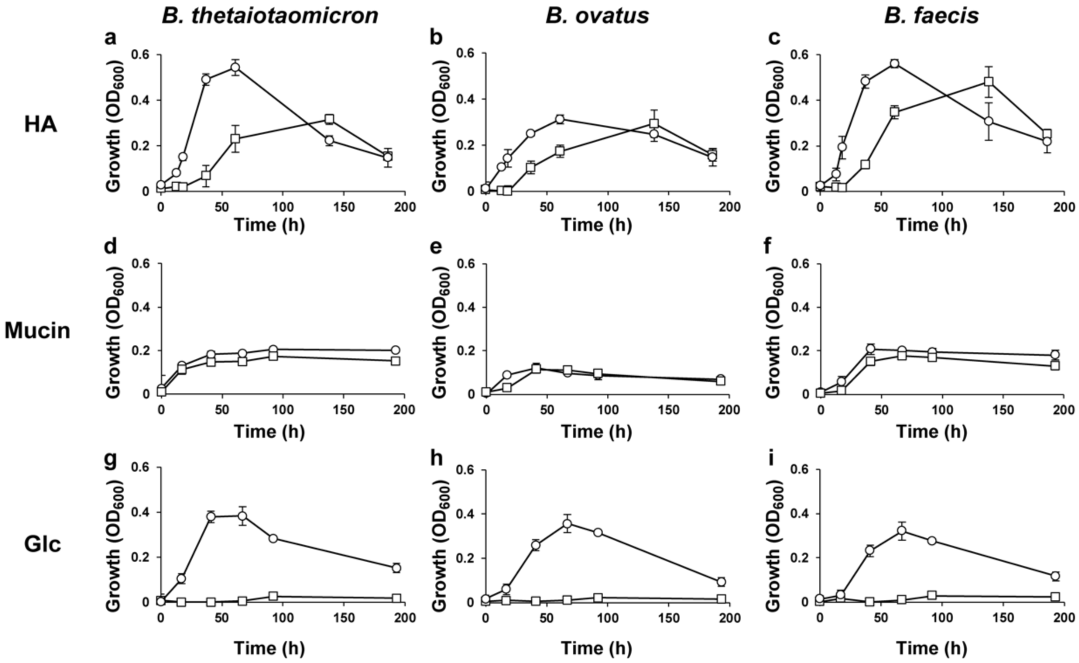


**Figure S7. *Bacteroides* species assimilate GAG (HA) or mucin as a sole carbon/nitrogen source.** Left, *B. thetaiotaomicron*; center, *B. ovatus*; right, *B. faecis*. The open circle and the square represent media with and without ammonium sulfate, respectively. Error bars show standard deviations (n = 3).

**Table S1. Summary of degradation assimilation of GAGs and mucin by *Bacteroides* species.**

| | GAG degradation | | | GAG assimilation | | | Mucin assimilation |
|---|---|---|---|---|---|---|---|
| | CSC | HA | HP | CSC | HA | HP | Mucin |
| *B. acidifaciens* | n.t. | n.t. | n.t. | n.t. | n.t. | n.t. | — |
| *B. barnesiae* | — | — | — | + | + | — | — |
| *B. caccae* | + | + | — | + | + | — | + |
| *B. cellulosilyticus* | + | — | — | + | — | — | — |
| *B. clarus* | + | + | + | + | + | + | + |
| *B. coagulans* | n.t. | n.t. | n.t. | + | + | — | n.t. |
| *B. coprocola* | — | — | — | + | + | — | — |
| *B. coprophilus* | — | — | — | + | + | — | — |
| *B. coprosuis* | — | — | + | — | — | — | + |
| *B. dorei* | — | — | — | — | — | — | — |
| *B. eggerthii* | + | — | + | + | — | + | — |
| *B. faecichinchillae* | — | — | — | — | — | — | — |
| *B. faecis* | + | + | + | + | + | + | + |
| *B. finegoldii* | + | + | + | + | + | + | + |
| *B. fluxus* | + | — | — | + | + | + | — |
| *B. fragilis* | — | — | — | — | — | — | + |
| *B. gallinarum* | + | — | — | + | + | + | + |
| *B. helcogenes* | — | — | — | — | — | — | + |
| *B. intestinalis* | + | + | + | + | — | + | + |
| *B. massiliensis* | + | — | — | + | + | + | — |
| *B. nordii* | — | + | — | + | + | + | + |
| *B. oleiciplenus* | + | — | — | + | — | — | — |
| *B. ovatus* | + | + | + | + | + | + | + |
| *B. paurosaccharolyticus* | + | + | — | — | — | — | n.t. |
| *B. salanitronis* | — | — | — | — | — | — | — |
| *B. stercoris* | + | + | + | + | — | + | + |
| *B. thetaiotaomicron* | + | + | + | + | + | + | + |
| *B. uniformis* | — | — | — | — | — | — | + |
| *B. vulgatus* | — | — | — | — | — | — | — |
| *B. xylanisolvens* | — | + | — | + | + | + | + |

n.t., not tested; +, positive in degradation or assimilation; —, negative in degradation or assimilation. Results (9) regarding GAG degradation are included.

**Table S2. Detection of *Bacteroides* PL8 and PL12 lyase genes in human feces.**

| | All bacteria 16S rDNA | *Bacteroides* PL8 lyase | *Bacteroides* PL12 lyase | Detection ratio (%)[a] |
|---|---|---|---|---|
| Sample A | 6.23 x $10^{11}$ | 1.47 x $10^{9}$ | 3.90 x $10^{9}$ | 6.03 |
| Sample B | 1.09 x $10^{11}$ | 6.45 x $10^{7}$ | 6.75 x $10^{7}$ | 0.85 |
| Sample C | 9.81 x$10^{10}$ | 6.11 x $10^{7}$ | 3.88 x $10^{8}$ | 3.20 |

[a] Detection ratio of PL8 and PL12 lyase genes was estimated based on the 7 copies of 16S rDNA in every bacterial cell and each 1 copy of PL8 and PL12 lyase genes in each *Bacteroides* cell. Mode of copies of 16S rDNA in superkingdom bacteria is shown to be 7 in The Ribosomal RNA Database (https://rrndb.umms.med.umich.edu).

**Table S3. Estimated essential amino acids which are produced by *Bacteroides* species in the human gut.**

| | *B. thetaiotaomicron* (mg)[a] | *B. ovatus* (mg)[a] | Recommended daily intake (mg)[b] |
|---|---|---|---|
| His | n.d. | 192 | 600 |
| Ile | 909 | 1,263 | 1,200 |
| Leu | 747 | 1,199 | 2,340 |
| Lys | n.d. | 120 | 1,800 |
| Met | 268 | 397 | 900 |
| Phe | 316 | 371 | 1,500 |
| Thr | 223 | 241 | 900 |
| Trp | n.d. | 153 | 240 |
| Val | 1,637 | 2,413 | 1,560 |

[a]The amount of essential amino acids produced by *Bacteroides* species was estimated using the following conditions. (i) The weight of gut microbiota is about 1.5 kg. (ii) *Bacteroides* species are dominant (~50% of total) in the human gut. (iii) Cells (2 g) of *Bacteroides* species were acquired from the GAG minimal medium (1,000 ml). n.d., not detected.

[b]The amount of essential amino acids per human adult (60 kg body weight) is recommended by the WHO.
